# A Benchmark of Computational CRISPR-Cas9 Guide Design Methods

**DOI:** 10.1101/498782

**Authors:** Jacob Bradford, Dimitri Perrin

## Abstract

The popularity of CRISPR-based gene editing has resulted in an abundance of tools to design CRISPR-Cas9 guides. This is also driven by the fact that designing highly specific and efficient guides is a crucial, but not trivial, task in using CRISPR for gene editing. Here, we thoroughly analyse the performance of 17 design tools. They are evaluated based on runtime performance, compute requirements, and guides generated. To achieve this, we implemented a method for auditing system resources while a given tool executes, and tested each tool on datasets of increasing size, derived from the mouse genome. We found that only five tools had a computational performance that would allow them to analyse an entire genome in a reasonable time, and without exhausting computing resources. There was wide variation in the guides identified, with some tools reporting every possible guide while others filtered for predicted efficiency. Some tools also failed to exclude guides that would target multiple positions in the genome. We also considered a collection of over a thousand guides for which experimental data is available. For the tools that attempt to filter based on efficiency, 65% to 85% of the guides they reported were experimentally found to be efficient, but with limited overlap in the sets produced by different tools. Our results show that CRISPR-Cas9 guide design tools need further work in order to achieve rapid whole-genome analysis and that improvements in guide design will likely require combining multiple approaches.

## 1 Introduction

Wild-type CRISPR (Clustered Regularly Interspaced Short Palindromic Repeats) is found in archaea and bacteria acting as an adaptable immune system [1]. CRISPR is able to provide a method of immunity via three steps [2]: (i) a DNA snippet from an invading phage is obtained and stored within the CRISPR array, making a memory of past viral infection; (ii) the CRISPR region is expressed and matured to produce duplicates of previously obtained DNA snippets (or *guides*); (iii) in the case of *S. pyogenes* Cas9 (SpCas9), a guide binds with a SpCas9 nuclease to enable site-specific cleavage due to guide-DNA homology. This last step provides immunity to the host cell and is also the mechanism for CRISPR to be used in a genome engineering context, where a synthetic guide is supplied. CRISPR-based systems have been used for a number of such applications [3–5]. However, guide design is not trivial, as the efficacy and specificity of guides are crucial factors of utmost importance. For this reason, computational techniques are employed to identify and evaluate candidate CRISPR-Cas9 guides.

Here, we analyse 17 CRISPR-Cas9 guide design methods to evaluate whether they are adequate for rapid whole-genome analysis, and potentially whether combining approaches would achieve a solution of better quality. The available tools can be categorised based on algorithmic approach (i.e. procedural or via models trained using experimental data); however, when constructing the tool-set, we considered various factors: (i) whether the source code was easily obtained, (ii) is the installation process straight-forward, (iii) the tool simply does not provide a wrapper for performing a regular expression and (iv) the guide length and PAM sequence be customised. In our analysis, we consider not only their output (i.e. which targets they identified), but also their ability to process whole genomes in a reasonable time. This is especially important for large genomes, such as some flowering plants. It is also a crucial feature for some applications such as studies of complex pathways or functions that require targeting multiple genes (e.g. sleep [6–8]) or producing whole-genome maps [9].

## 2 Tool Review

We selected 17 guide design tools that are released under an open-source license and report candidate guides for the *Streptococcus pyogenes*-Cas9 (SpCas9) nuclease; these tools are listed in Table 1.

**Table 1.**
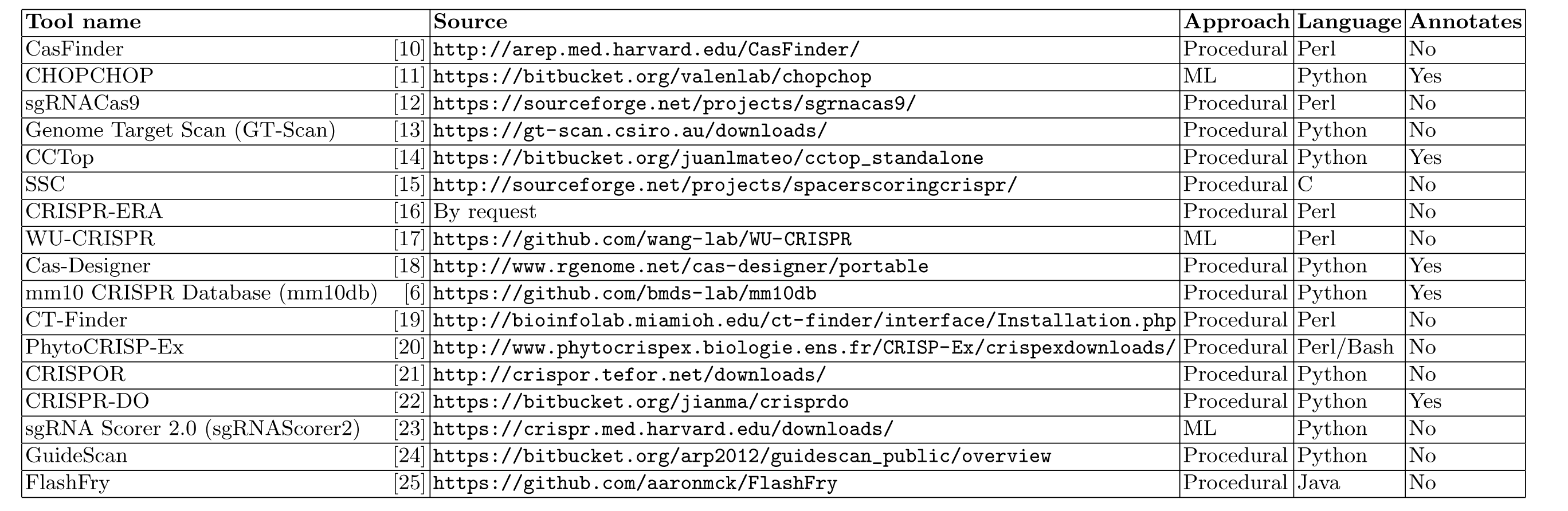
The seventeen selected tools analysed in this study; chronologically ordered. *ML*: machine learning

Python (n=9) and Perl (n=5) are the most common programming languages, with CT-Finder and CRISPOR also implementing web-based interfaces via PHP (complemented by JavaScript, CSS, HTML, etc.). To improve run time, mm10db implements some of its components in C. SSC and FlashFry are implemented in C and Java, respectively. PhytoCRISP-Ex is a Perl-implemented tool, however, extensively utilises Linux bash commands for pre-processing. Configuration of tools is most commonly achieved via command-flags, however, tools such as Cas-Designer and CasFinder are configured via a text file. CHOPCHOP is configured via global variables in the main script file.

SciPy [26] (inc. Numpy) and BioPython [27] were common packages utilised by the Python-based tools. CHOPCHOP and WU-CRISPR use SVMlight [28] and LibSVM [29], respectively; both being C implementations of support vector machines (SVM). Similarly, sgRNAScorer2 utilsies the SVM module from the SciKit-learn package [30]. The authors supplied models for the 293T cell line and the Hg19 and mm10 genomes. sgRNAScorer2 implements a SVM via the SkLearn package and supplies a model trained by high- and low-activity guides. CHOPCHOP, GT-Scan, CRISPR-ERA, CCTop and CasFinder use Bowtie [31] for off-targeting; similarly to mm10db and CT-Finder with Bowtie2 [32]; while CRISPOR and CRISPR-DO utilise the Burrow-Wheelers Algorithm (BWA) [33]. GuideScan does not depend on external tools for off-targeting, and instead implements a trie structure for designing guides with greater specificity [24]. FlashFry benefits from its guide-to-genome aggregation method which is able to identify off-target sites in a single pass of its database. This achieved greater performance for FlashFry in comparison to BWA, as the number of mismatches and number of candidate guides increases [25]. Interestingly, [34] identified that Bowtie2 lacks the ability to rapidly identify all off-target sites with greater than two mismatches and that Cas-OFFinder [35] is a more suitable solution, however is more time expensive.

CHOPCHOP, Cas-Designer, mm10db, CCTop and CRISPR-DO provide ways to specify annotation files. For each of these tools, we provided the appropriate annotation in each test. CHOPCHOP utilises the annotations to indicate which gene or exon a candidate guide targets, and to allow the user to restrict the search region to particular genes. Cas-Designer utilises a custom-format annotation file, which describes the start and end positions of each exon on a given genome. This is used for designing candidate guides which specifically target exon regions. mm10db requires an annotation file in the UCSC RefGene format in order to generate a file containing sequences for all exon regions. CCTop utilises annotations for evaluating guides based on their distance to the closest exon, and for passing results to the UCSC genome browser as a custom track.

Some biological rules are shared across tools, such as: avoiding poly-thymine sequences [36], rejecting guides with inappropriate GC-content [37], counting and possibly considering the position of single-nucleotide polymorphisms (SNPs) and considering the second-structure formation. Most tools report all targets that have been identified (sometimes with a score for specificity and/or efficiency) and rely on the user to determine whether a guide is appropriate for use, while mm10db actively filters guides and only reports ‘accepted’ ones (but ‘rejected’ targets, and reason for rejection, are still available in a separate file). However, some tools (sgRNAScorer2 and WU-CRISPR) do not implement any of these rules through procedural-styled programming but instead, utilise machine learning models trained from experimental data. Furthermore, due to the age of some tools and the rapid growth of CRISPR-related research, the specifics of these procedural rules vary. For example, early studies describe the 10-12 base pairs adjacent to the PAM (the *seed* region) are more significant than the remainder of the guide [1, 38], however, recent research contradicts this and suggest that one-to-five base pairs proximal to the PAM are more likely to be of significance [39]. When evaluating tools, these may be factors that researchers would want to consider.

## 3 Method

Given the range of rules and implementations, it is crucial to benchmark guide design tools and compare their performance. In this section, we describe how each tool is evaluated based on compute requirements, features and output.

### 3.1 Data Preparation

The initial data from our benchmarking is based on the *GRCm38/mm10* mouse genome assembly, available via the University of California, Santa Cruz (UCSC). We downloaded chr19, and extracted three datasets of increasing length: 500k, 1m, and 5m nucleotides, all starting from position 10,000,000. These datasets, and the whole chromosome, are used for testing.

For each of these four configurations, we created all the files required by any of the tools: custom annotation file (derived from the *refGene* table available via UCSC), 2bit compression file, Bowtie and Bowtie2 indexes, and Burrows-Wheeler Aligner file.

To complement these datasets, we have also used a collection of guides for which experimental data is available [15]. This collection contains 1169 guides which were used in a screening experiment, with 731 deemed to be ‘efficient’ based on analysis of the gene knock-outs. We constructed an artificial sequence that contains these guides, interspaced by 50 Ns to ensure that unexpected overlapping targets cannot be detected. As before, we generated all the supporting files required for this input.

The tools were not optimised for a specific organism, and the choice of mm10 for the initial tests, or of data from human cell lines for the experimental validation, should not impact the results

Our datasets are available in the Supplementary Materials.

### 3.2 Performance Benchmarking

Our *Software Benchmarking Script* (SBS) tool is implemented in Python 2.7, and uses the Processes and System Utilisation (PSUtil, version >= 5.4.4) module for process-specific monitoring of system resources. When launched, the user is required to pass a bash command and an output directory via command-line flags. The audit routine begins after the bash command is executed. The parent process and all descendants are monitored at each polling event (PE).

The current wall-time, CPU and memory usage, disk interaction (DIO), number of threads and number of children are recorded. Wall-time was preferred to CPU-time as it is the human-perceived completion time. SBS reports the instantaneous resident set size (RSS) usage and virtual memory usage. DIO includes both the number and size of read/write operations. At each PE, an aggregate of the parent and child data is calculated and written to file. Additionally, the bash commands which launched each child process are logged.

The PE routine continues until the parent process ends or 72 hours is exceeded. This limit was imposed as we aimed to discuss tools with potential for whole-genome analysis, and those that cannot analyse chr19 (2.25% of mm10 total size) within this limit are deemed inappropriate for the task.

All tests were performed on a Linux workstation with Intel Core i7-5960X (3.0 GHz), 32 GB RAM, 32 GB allocated swap space, and Samsung PM87 SSD. We used Python v2.7 and Perl v5.22.1. This machine exceeds the specifications of some workstations, however, it is expected that a user would require a similar machine or better in order to achieve whole-genome analysis.

SBS is available on GitHub at https://github.com/jakeb1996/SBS.

### 3.3 Output Normalisation & Comparison

Each tool had its own output format, so we normalised the results as: tool name, candidate guide sequence, start position, end position and strand (written to file in the CSV format). During this process, the start and end values were aligned with one-based positioning, as per the UCSC datasets. An ellipsis was concatenated to guides which were lacking a PAM sequence.

To determine which tools shared common guides, a script aggregated all non-duplicate guides and recorded which tool produced subsequent occurrences of the guide. A guide was considered a duplicate with a previously observed guide when the 3’ positions were equal and when they were reported to target the same DNA strand. A separate script analysed each normalised guide to determine whether it targeted a gene coding region (based on the UCSC annotation data).

## 4 Results

Each tool was tested sequentially on increasing input sizes. If a tool exceeded the time limit or was terminated by the operating system, it did not proceed to larger datasets. CT-Finder, GT-Scan and Cas-Designer reached the time limit on the ‘full’ dataset; sgRNAScorer2 and PhytoCRISP-Ex reached it for the ‘5m’ dataset; FlashFry reached it for the ‘1m’ dataset. The out-of-memory killer terminated CHOPCHOP and CRISPOR on the ‘full’ dataset. The 30 Gb of allocated memory for the Java virtual machine was saturated when running FlashFry. SSC only successfully executed on the 500k dataset; testing on datasets of a larger size resulted in the tool returning an *processing failed* error message. WU-CRISPR was unable to analyse the full dataset as the unmodified UCSC mm10 chromosome 19 file contained N’s, which the tool did not accept.

### 4.1 Output

A similar set of candidate guides was generated by each tool, however, the start and end positions of identical guides often differed by up to 4 positions. This was seen due to some tools truncating the PAM sequence, or where zero-based positioning was used. An ellipsis was concatenated during normalisation for those that truncated it. All guides were aligned with one-based positioning as per the UCSC datasets, however, it is noted that Ensembl and Bowtie adhere to a zero-based system, explaining its usage in some tools. Many tools produced output using the comma- or tab-separated values (CSV, TSV, respectively) formats (n=16), both of which are compatible with software such as Excel. Some of these tools did not provide a header line to indicate the meaning of each column. GT-Scan differed as it produced an SQLite database containing a table of all candidate guides. Tests which were terminated did not produce output as the write-to-file routines had not been reached prior to termination.

### 4.2 Output Consensus

Here, we discuss the consensus between tools for the datasets derived from chr19. We focus on the ‘500k’ dataset, as this was the only dataset where each tool successfully completed a test. Similar results were observed on the larger datasets, for the tools that still managed to produce an output.

We define the consensus *C_i,j_* between tool *i* and tool *j* as the proportion of guides produced by *i* that are also produced by *j*. Note that the value is not symmetric: while the intersection between the two set of guides is unchanged, the denominator for *C_i,j_* and *C_j,i_* is the number of guides produced by *i* and by *j*, respectively.

The consensus matrix *C* is shown in Fig. 1A, and highlights that most tools do not filter targets. They report all possible targets, sometimes with a score. This leads to a high consensus between methods, typically 95% or more. The CHOPCHOP method, on the other hand, checks that a guanine is present at the twentieth nucleotide. If this is enforced, it reports only about a quarter of the targets that other tools produce. CRISPR-DO removes those that contain polythymine sequences and those with undesirable off-target effects. PhytoCRISP-Ex only considers as potential targets the guides that satisfy two rules: (i) they have at most two off-target sites with only one mistmatch anywhere else in the input genome, and (ii) their seed region (last 15 bases including the PAM sequence) is unique. These potential guides are then checked for the presence of restriction sites for pre-selected, common restriction enzymes. The consensus of WU-CRISPR with other tools was a maximum of 27.2%. This was due to the tool applying pre-filters to eliminate guides before being evaluated by the SVM model. Some of the pre-filters included: folding and binding properties and position specific guide contents. Following this, the tool only reports guides which match the model.

**Fig. 1.**
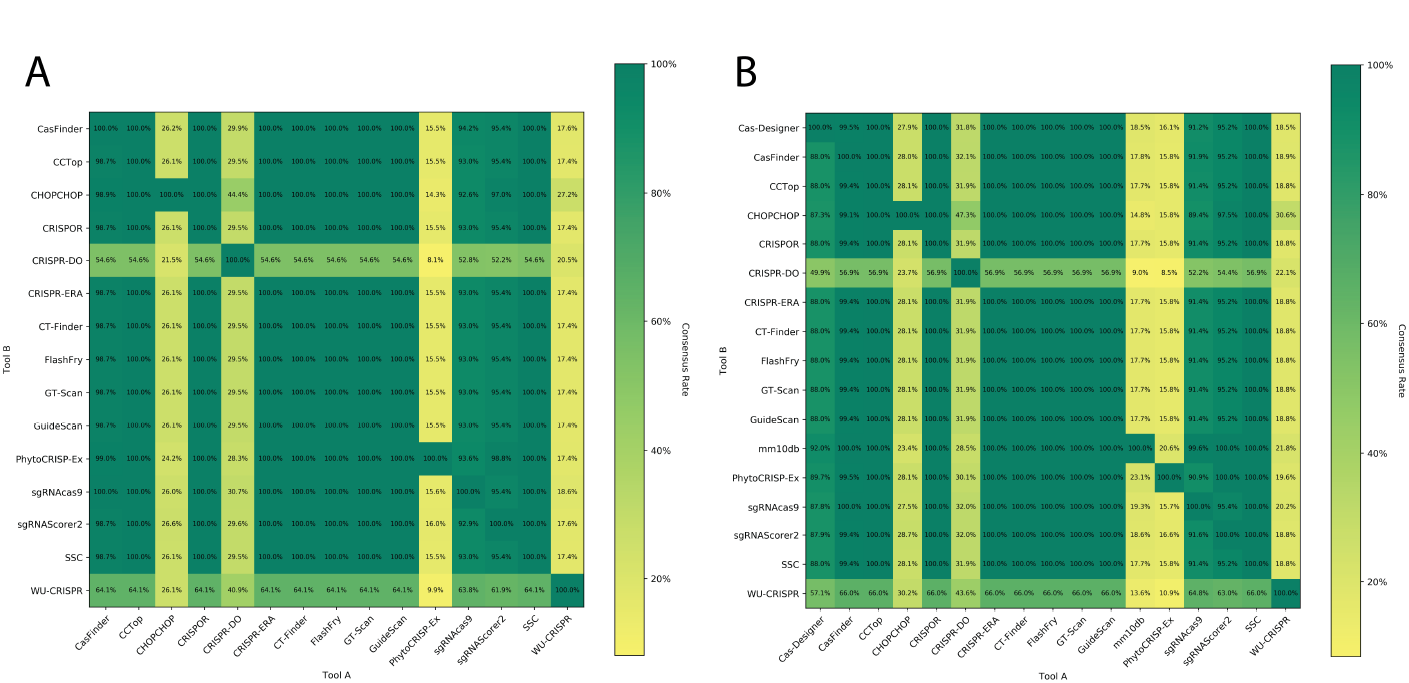
Consensus matrix (‘500k’ dataset). (A) Consensus between tools that target the entire input dataset. (B) Consensus between all tools, for coding regions only.

Some tools focus (or give the option to focus) on identifying targets located on exons. Fig. 1B shows the consensus matrix for these regions, across all tools. As before, there is a high consensus between tools that do not reject targets. For tools that more strictly select which targets to report (CHOPCHOP and mm10db), the consensus is again around 15-30%. Interestingly, the consensus between CHOPCHOP and mm10db is low. This highlights that tools are using different selection criteria, and this leads to the identification of distinct targets.

We also analysed the reasons provided by mm10db for rejecting guides. It is worth noting its filters are applied sequentially. For instance, a guide rejected for multiple exact matches in the genome might have also been rejected for its GC content if it had made it to that filter. The results are still informative. The main reasons were high GC-content (approx. 62%), a poor secondary structure or energy (19%), and multiple exact matches (16%). Off-target scoring is the last step so, despite being strict, it necessarily contributes to a small fraction of the rejections. Here, this is made even smaller (< 0.05%) because the small input size limits the risk of having a large number of similar off-target sites.

From Fig. 1, we know that many guides proposed by other tools are rejected by mm10db. For each tool, we also explored the rejection reasons given by mm10db. This is summarised in Fig. 2 for the ‘500k’ dataset. Rejection due to GC content is the most significant reason, followed by secondary structure or energy.

**Fig. 2.**
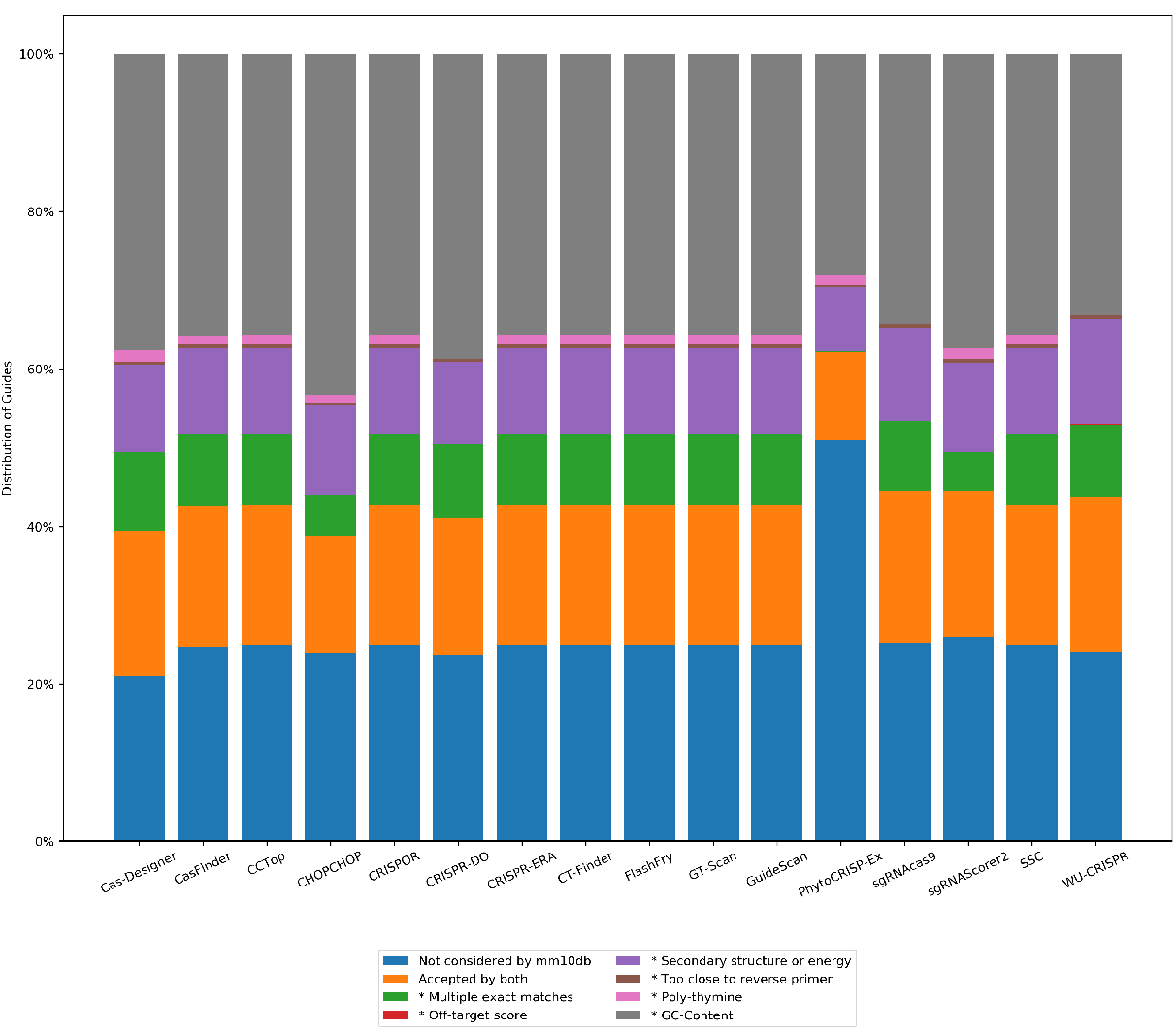
Reasons for rejection. This plot shows, for each tool, the proportion of accepted guides that have been rejected by mm10db, and the reason for doing so. An asterisk (*) indicates the category relates to a rule implemented in mm10db for rejecting guides.

Importantly, a number of guides (on average around 10%) proposed by these tools are rejected by mm10db due to multiple exact matches in the genome. For instance, guide CTCCTGAGTGCTGGGATTAAAGG is reported by 10 tools, despite targeting two distinct genes on chr19 alone: Prpf19 at chr19:10,907,131-10,907,153 and Gm10143 at chr19:10,201,455-10,201,477. Ideally, this guide would not appear in the output from a tool due to its lack of specificity: there is no practical application where such a guide would be useful. Furthermore, none of the tools that deal with specificity by reporting the number of sites with *k* mismatches (*k* ≤ 5) actually reported the correct number of off-targets for this guide. Cas-Designer and GT-Scan reported zero perfect matches. CHOPCHOP reported 50 perfect matches (Bowtie was used by CHOPCHOP for this task, however, was limited to report 50 alignments). Aligning this guide on the full chr19 using Bowtie (*-v 0 -a*), we found 241 perfect matches. However, to the contrary, PhytoCRISP-Ex did not report any guides which had multiple matches in the genome. This is because, as mm10db, it is strict about filtering out guides that may have off-target partial matches.

Fig. 2 also shows that, for tools that report all guides, roughly a quarter of their output has not been considered (i.e. neither accepted nor rejected) by mm10db. This is due to the regular expression it is using to extract candidates ([*ACG*][*ACGT*]^20^*GG*), which exclude candidates starting with a T, while other tools do not.

### 4.3 Experimental Dataset

We then considered the performance of tools that either filter targets and provide a score to predict efficiency, as seen in Figure 3. We evaluated them on the artificial genome constructed with experimentally validated guides.

**Fig. 3.**
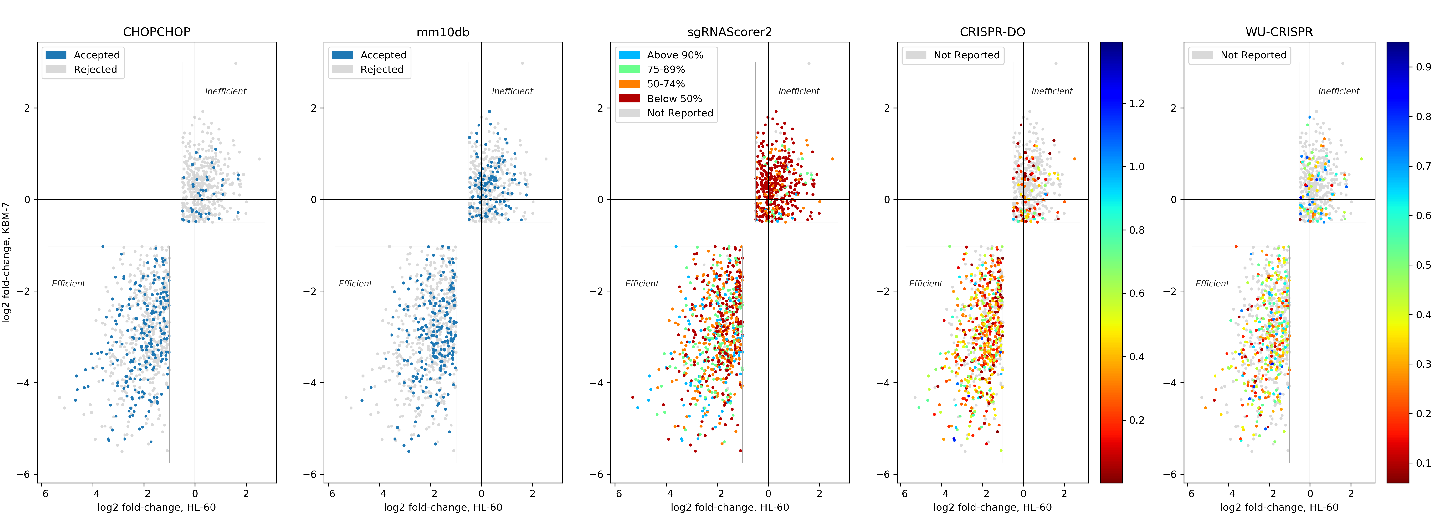
Tool performance on experimentally validated guides. For CHOPCHOP and mm10db, colours indicate the whether a guide was accepted by the tool. For sgRNAScorer2, CRISPR-DO and WU-CRISPR, colours represent the score distribution within this dataset.

CHOPCHOP reported 273 guides, 84.3% of which were ‘efficient’, and mm10db reported 330, 65.2% of which were ‘efficient’. Incorporating CHOPCHOP’s rule of having a G at position 20 into mm10db would bring its proportion of ‘efficient’ guides to 84.4%. The results also highlight the value in considering multiple selection approaches: only 25.1% of the ‘efficient’ guides identified by mm10db have also been selected CHOPCHOP.

The sgRNAScorer2 user manual states that scores should be used in a *relative* manner as opposed to *absolute*: the higher the score, the better the predicted activity, but there is no threshold to classify between predicted efficiency and inefficiency. The 50th, 75th and 90th percentiles used in Fig. 3 are therefore for visualisation only. Instead, we extracted every pair (*a,b*) of guides where *a* was experimentally considered ‘efficient’ and *b* ‘inefficient’, and checked whether *a* had a higher predicted activity than *b*. This was true for 76.8% of them.

CRISPR-DO and WU-CRISPR did not report all the guides provided in the artificial genome, but for those that they did, an absolute score was reported. These tools reported 590 (84.6% efficient) and 402 (77.9% efficient) of guides, respectively. CRISPR-DO’s results reflect that it has the highest rate of reporting guides which were deemed experimentally efficient.

For the five tools considered, there is a very low consensus, at 3.6% (26 of 726). While it is not possible to generalise too much from a single dataset, the results suggest that all five tools have the ability to detect some efficient guides, but also that they are not completely precise and still have low recall. Improved methods are needed.

### 4.4 Computational Performance Analysis

Even though it is often overlooked when methods are initially presented, their computational performance is an important consideration. There are a number of applications of CRISPR where it can be crucial: large input genomes, time constraints to obtain the results, large number of non-reference genomes to analyse, etc.

In terms of time requirements, our hypothesis was that any efficient scoring or filtering would scale linearly with the number of targets to process (since they are individually assessed), and therefore almost linearly with the input genome size. However, for the specificity analysis, each guide needs to be assessed against any other candidates, which could result in quadratic growth. Another possible limitation is the memory requirement of the tools.

Our results are shown in Table 2. Only five tools successfully completed the four tests: CasFinder, CRISPR-ERA, mm10db, GuideScan and CRISPR-DO. Three of these reported all possible targets in Section 4.2, with no scoring on predicted efficiency, and were therefore expected to be fast. On the other, mm10db runs candidate guides through a number of filters.

**Table 2.**
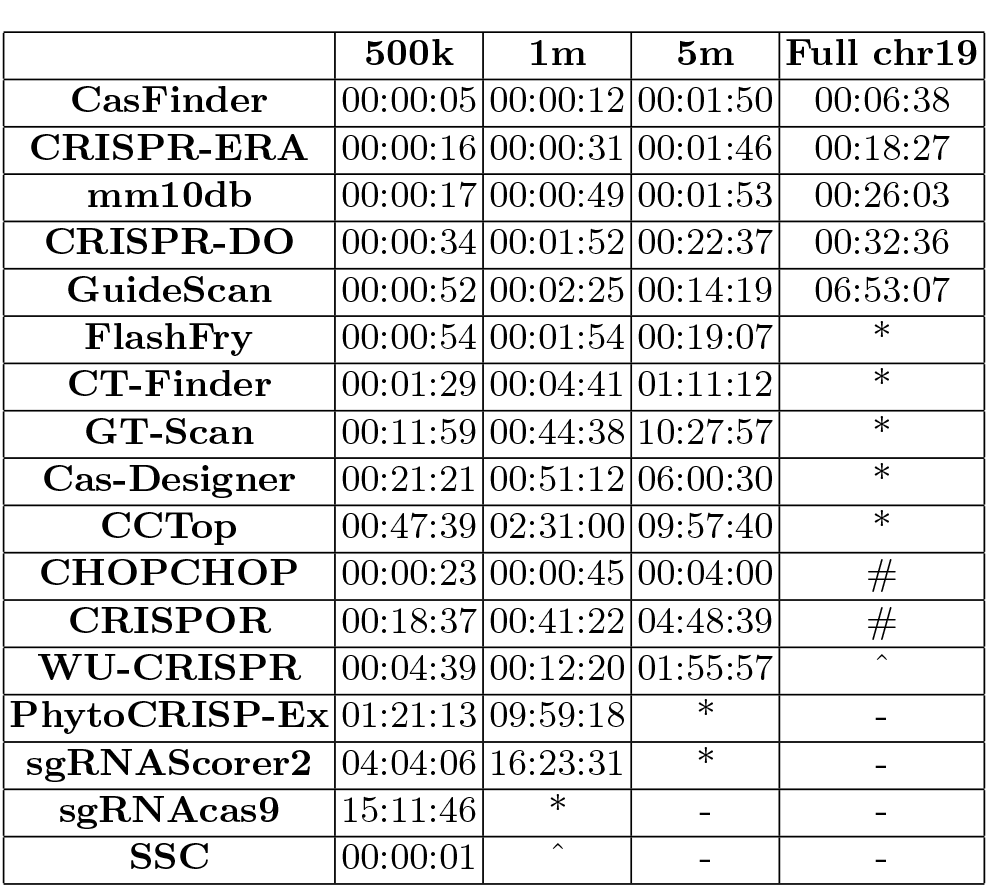
Run time (hh:mm:ss) for each tool and dataset size. Some tools exceeded the 72-hour limit (*), or were terminated by the the operating system’s out-of-memory killer (#). Any tool failing a test was not considered for larger input sizes (-). WU-CRISPR failed on ‘full’ due to the tool rejecting inputs containing N, and SSC could not process inputs larger than 500,000 bases in length (^).

The two tools that provide filtering or scoring based on predicted efficiency were not able to process the entire chromosome. CHOPCHOP saturated both physical memory and allocated swap space (as monitored by SBS), resulting in the out-of-memory killer (OOMK) terminating its run on the full chromosome 19 (approx. 2% of the mouse genome). sgRNAScorer2 was extremely slow, taking more 4 hours to process the 500k dataset (when the other tools took between 5 seconds and 21 minutes). It did not manage to process the 5m dataset in less than 3 days. FlashFry was performing well in the tests below ‘full’, however, the Java Virtual Machine was exhausted of memory when completing the final test.

Four other tools failed to satisfy the time constraint, all on the full chromosome 19 (Cas-Designer, CT-Finder, GT-Scan and CCTop). CRISPOR, had memory problems similar to those of CHOPCHOP and was terminated on that same dataset.

Fig. 4 summarises these results and takes into account that some tools only considered exons. We normalised the computation time of each tool by the total length of the sequences it considered, thus obtaining a measure of ‘effective basepairs per second’ (ebp/s). Three groups are visible: those tools with a measurable run-time for all four tests, (ii) those tools with a measurable run-time for three of the tests and (iii) those that only successfully completed one test.

**Fig. 4.**
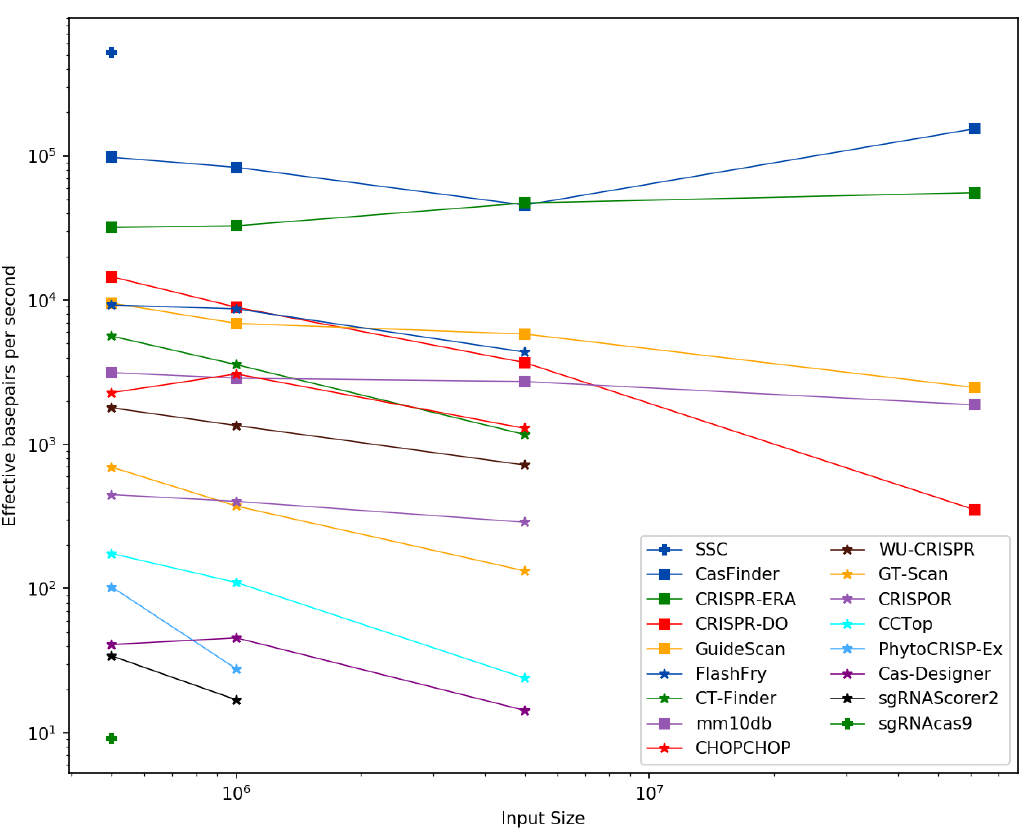
Normalised processing speed (in effective basepairs per second), taking into account the total length of the sequences considered by any given tool. Three performance groups have been identified, and are shown with different symbols for their markers.

The first group contains the two most rapid tools that were able to complete more than one test: CasFinder and CRISPR-ERA. The performance of other tools in this group is significantly poorer for all tests. However, given the results from Section 4.4, the immediate value of mm10db, and the potential for CHOPCHOP (either as a reimplemented, memory-conscious tool or for some its features as added components to mm10db) presents great opportunity in the development of future guide design tools. sgRNACas9 and sgRNAScorer2 remain the slowest tools overall, both requiring exorbitant time to complete analysis on the smallest dataset.

As part of our analysis we also considered the use of multi-threading. CHOP-CHOP, GuideScan, mm10db, FlashFry and sgRNAScorer2 are the only tools which have implemented multi-threaded routines. CHOPCHOP and GuideScan do not allow the user to specify the number of threads to spawn, but instead, spawns according to the number of CPU cores. mm10db provides command-flags to specify the number of threads for itself and Bowtie; we specified 128 and 8 threads, respectively. It is the only tool that enables multi-threaded mode for Bowtie. Cas-Finder, CT-Finder, GT-Scan and CHOPCHOP utilise Bowtie in single-threaded mode.

SBS monitored the physical memory and allocated swap space usage. CRISPOR and CHOPCHOP saturated both of these memory spaces, resulting in the out-of-memory killer (OOMK) terminating each on the full dataset test. Cas-Designer saturated the swap space, however did not trigger the OOMK in the same test.

## 5 Conclusion

A large number of tools have been proposed to assist in the design of CRISPR-Cas9 guides. Many have them have been successfully used experimentally: Cas-Finder has been used for designing guides from the human genome to target specific genes [40], CRISPR-ERA has been used for designing guides to target the HIV-1 viral genome [41] and mm10db has been used for whole-genome analysis of the mouse genome [6].

However, there has been little comparison of their performance. This paper address this gap by benchmarking 17 tools in terms of their computational behaviour as well as the guides they produce. Our results show that only five tools (CasFinder, CRISPR-ERA, CRISPR-DO, GuideScan and mm10db) can claim to analyse large inputs rapidly and would be readily available to process entire genomes, especially larger ones.

Most tools are very selective in designing guides and therefore have a high consensus between them. Five tools provide clear predictions of guide efficiency (sgRNAScorer2, CHOPCHOP, mm10db, CRISPR-DO and WU-CRISPR), and we assessed their performance on a collection of experimentally validated guides. All five tools produced a majority of efficient guides, but interestingly there was only limited overlap between them.

Taken together, all these results highlight the need for further refinement of CRISPR-Cas9 guide design tools, and provide a clear direction for future work in optimising computational performance and combining multiple design approaches.

